# A model of flux regulation in the cholesterol biosynthesis pathway: Immune mediated graduated flux reduction versus statin-like led stepped flux reduction

**DOI:** 10.1101/000380

**Authors:** Steven Watterson, Maria Luisa Guerriero, Mathieu Blanc, Alexander Mazein, Laurence Loewe, Kevin A Robertson, Holly Gibbs, Guanghou Shui, Markus R Wenk, Jane Hillston, Peter Ghazal

**Author notes:** PRESENT ADDRESSES:-Systems Biology Ireland, Conway Institute, University College Dublin Belfield, Dublin 4, Ireland (Maria Luisa Guerriero). Wisconsin Institute for Discovery, 330 North Orchard Street, University of Wisconsin-Madison, Madison, WI 53715, USA. Tissue Microscopy Laboratory, Department of Biomedical Engineering, 337 Zachry Engineering Center, 3120 Texas A&M University, College Station, TX 77843, USA.

## Abstract

**Graphical Abstract:** 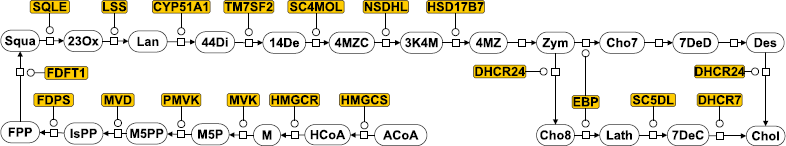

**Highlights:**

- We model the cholesterol biosynthesis pathway and its regulation
- The innate immune response leads to a suppression of flux through the pathway
- Statin inhibitors show a different mode of suppression to the immune response
- Statin inhibitor suppression is less robust and less specific than immune suppression

**Asbtract:** The cholesterol biosynthesis pathway has recently been shown to play an important role in the innate immune response to viral infection with host protection occurring through a coordinate down regulation of the enzymes catalyzing each metabolic step. In contrast, statin based drugs, which form the principle pharmaceutical agents for decreasing the activity of this pathway, target a single enzyme. Here, we build an ordinary differential equation model of the cholesterol biosynthesis pathway in order to investigate how the two regulatory strategies impact upon the behaviour of the pathway. We employ a modest set of assumptions: that the pathway operates away from saturation, that each metabolite is involved in multiple cellular interactions and that mRNA levels reflect enzyme concentrations. Using data taken from primary bone marrow derived macrophage cells infected with murine cytomegalovirus infection or treated with IFN*γ*, we show that, under these assumptions, coordinate down regulation of enzyme activity imparts a graduated reduction in flux along the pathway. In contrast, modelling a statin-like treatment that achieves the same degree of down-regulation in cholesterol production, we show that this delivers a step change in flux along the pathway. The graduated reduction mediated by physiological coordinate regulation of multiple enzymes supports a mechanism that allows a greater level of specificity, altering cholesterol levels with less impact upon interactions branching from the pathway, than pharmacological step reductions. We argue that coordinate regulation is likely to show a long-term evolutionary advantage over single enzyme regulation. Finally, the results from our models have implications for future pharmaceutical therapies intended to target cholesterol production with greater specificity and fewer off target effects, suggesting that this can be achieved by mimicking the coordinated down-regulation observed in immunological responses.

## 1 Introduction

Cholesterol is central to a diverse range of cellular functions, including membrane development and maintenance [1], lipid raft formation and vesicular transport [2], steroid hormone synthesis [3], neurological development [4], and oxysterol and vitamin D synthesis [5]. Recently, the cholesterol metabolism has been shown to have an important role in host-pathogen interactions. It has been documented to be perturbed in response to infection [6, 7] and, conversely, cholesterol and its associated metabolites have been shown to alter inflammatory mediators [8, 9].

Cholesterol synthesis is one step in a pathway of metabolic interactions that is subject to catalytic regulation [10] and evidence suggests that this pathway is critical to the optimal growth of a range of viruses and microbes including cytomegalovirus (CMV), Hepatitis C (HCV), HIV, Japanese Encephalitis (JEV), West Nile (WNV), Dengue (DENV), Measles viruses (MV), African Swine Fever Virus (ASFV), Mycobacteria and Salmonella [6, 7, 11, 12, 13, 14, 15, 16, 17, 18].

The cholesterol biosynthesis pathway itself comprises a sequence of metabolic interactions that occur across several organelles, starting with the processing of Acetyl-Coenzyme A (henceforth denoted ACoA - Supplementary section 1 for a list of all metabolite abbreviations) in the mitochondria and ending with cholesterol synthesis in the endoplasmic reticulum (***cite Mazein et al, same issue***)[**?**, 19, 20]. This pathway branches in the peroxisome and endoplasmic reticulum, into the sterol arm and the non-sterol arms (prenylation and dolichylation), the latter arms carrying flux away from the main sterol arm.

Coordinate transcriptional control of the enzymes of the cholesterol biosynthesis pathway is mediated by SREBP2 and feedback control occurs through regulation of SREBP2 transport. The SCAP:SREBP2 complex is ordinarily chaperoned to the Golgi complex where SREBP2 is cleaved, before it migrates to the nucleus to activate the suite of enzymes associated with the pathway. However, in the presence of relatively high concentrations of intracellular cholesterol or side-chain hydroxylated cholesterol, in particular 25-hydroxycholesterol, SCAP:SREBP2 is retained instead in the endoplasmic reticulum. Retaining SCAP:SREBP2 acts to down-regulate transcription of the enzymes acting on the pathway until ordinary levels of cholesterol and its derivatives have been restored. Hence, the pathway undergoes transcriptionally mediated regulation through changes to enzyme concentrations [19].

Recently, we reported a modest, but statistically significant, decrement in the concentrations of enzymes associated with cholesterol biosynthesis pathway, in response to both infection and interferon treatment in macrophages. This was observed at the transcriptional level and was shown to correlate with reduced protein concentrations [11]. This decrement was found to be part of the innate immune response, intended to suppress viral growth. However, the mechanism through which such changes decrease the activity of the cholesterol biosynthesis pathway is something that has yet to be fully elucidated in the published literature.

We have sought to investigate this mechanism of regulation, exploring the impact on flux that results from such enzyme decrements. This problem is experimentally challenging, but tractable with computational methods. Flux is the natural quantity to consider when studying metabolic pathway function and flux studies have been employed both theoretically [21, 22] and experimentally [23, 24]. The flux through the pathway describes the stoichiometrically adjusted rate of production of each metabolite and so captures whether and how the production rate of the metabolites affect each other. Ultimately, the final flux value in the pathway describes the rate of cholesterol synthesis. Metabolic Control Analysis (MCA) and Flux Balance Analysis (FBA) are two typical approaches to studying flux in a pathway system. However, MCA approaches focus on the effects of individual infinitesimal changes in enzyme activity rather than the compound effects of multiple finite changes and FBA, in its standard form, does not relate flux changes to substrate concentration changes. As a result, they are inappropriate for our study in which we validate the pathway model at the level of the substrate, implement multiple finite enzyme decrements and model the effects of chemical inhibition. We build an ordinary differential equation (ODE), dynamical model of the sterol pathway using Michaelis-Menten and mass action kinetics that incorporates additional interactions to represent the consumption of metabolites in non-sterol related processes. We demonstrate that multiple small decreases in enzyme activity can suppress the flux through the main cholesterol biosynthesis pathway. This suppression presents itself as a graduated reduction when the profile of flux is considered.

Cholesterol levels have also been demonstrated to be an important risk factor in cardiovascular disease [25, 26] and their control is an active area of research [27]. Current therapies involve the use of statins to competitively inhibit the enzyme HMGCR which is responsible for catalysis of the interaction transforming 3-hydroxy-3-methyl-glutaryl Coenzyme A (HCoA) to Mevalonate (M). However, the efficacy of such therapies is limited by drug toxicity and off target effects [28, 29, 30]. Here, we show that statin treatment regulates the flux through the pathway in a manner that is markedly different to that following infection. The metabolic interaction catalyzed by HMGCR is significantly upstream of cholesterol biosynthesis and we show that the impact of a statin-like treatment is to suppress flux throughout most of the pathway. This impacts significantly upon many of the metabolites upstream of cholesterol and upon the non-sterol arms, thereby incurring off-target effects. In contrast, because coordinate enzyme regulation leads to a graduated reduction in flux along the pathway, it has a less dramatic impact upon the branches upstream of cholesterol production.

This manuscript is organized as follows. In Section 2, we describe the experimental and mathematical methods employed to determine enzyme and metabolite levels in response to infection and IFN*γ* treatment and to model the pathway. In Subsection 2.1, we describe the experimental method and in subsections 2.2 and 2.3, we describe how the model was built, how the initial conditions were defined and how the model was used to simulate pathway activity. In Section 3, we present the results of using the model to study the flux through the pathway, with subsections 3.1, 3.2, 3.3 and 3.4 describing the validation of the model and the impact on the flux of the response to IFN*γ* treatment, to CMV infection and to statin intervention, respectively. In Section 4, we discuss these results, their relationship, their off-target effects and their implications for specific, targeted regulatory strategies. In Section 5, we summarize our results. Supplementary material in support of the results presented here is available online.

## 2 Materials & Methods

### 2.1 Experimental measurements

Enzyme levels were inferred from gene expression measurements of bone-marrow derived macrophage cells in two time course experiments, one in which cells were infected with murine cytomegalovirus (mCMV) and one in which cells were treated with IFN*γ*. Measurements were taken at half hour intervals for 12 hours using Agilent microarray platforms and at 24 hours for select members using QPCR. Agreement between mRNA expression and protein concentrations was validated by quantitative western blotting for selected members [11].

Intracellular cholesterol concentration was determined enzymatically using the Amplex-Red cholesterol assay kit (Molecular Probes) according to manufacturer recommendations. Briefly, cells were washed with 1 ml ice cold PBS and then lyzed in 200 *µ*l cold Lipid buffer containing 0.5M of potassium phosphate, pH 7.4, 0.25 mM cholic acid, and 0.5% triton X-100. Cell lysates were sonicated on ice with three 10-second pulses at high intensity. 20 *µ*l were then used to determine protein concentration using a standard BSA assay to normalize the protein concentration. For cholesterol measurement, 20 *µ*l of each sample were added to 80 *µ*l assay solution, which contained 300 *µ*M Amplex Red reagent, 2 U per ml HRP and 2 U per ml cholesterol oxidase, 0.1M of potassium phosphate, pH 7.4, 0.05mM cholic acid, and 0.1% triton X-100. After preincubation for 30 min at 37 *o*C under light exclusion conditions, fluorescence was measured using excitation at 530±2.5 nm and fluorescence detection at 590±2.5 nm with a Polarstar Optima Multifunction Microplate Reader (BMG Labtech, UK). The values were corrected from the background. The relative amount of free cholesterol to the mock treated samples was calculated using the manufacturer’s supplied standard curve.

For the measurement of metabolite concentration, an Agilent high performance liquid chromatography (HPLC) system coupled with an Applied Biosystem Triple Quadrupole/Ion Trap mass spectrometer (4000Qtrap) was used for quantification of individual polar lipids (phospholipids and sphingolipids). Electrospray ionization-based multiple reaction monitoring (MRM) transitions were set up for the quantitative analysis of various polar lipids. HPLC atmosphere chemical ionization (APCI)/MS were carried out for analysis of sterols [11].

### 2.2 Model Construction

Because regulation and feedback occur through transcriptional control of enzyme activity, we chose to model the impact of enzyme activity on the pathway flux. This obviated the need to explicitly consider SREBP2 mediated feedback in the pathway as any feedback would be accounted for in our measurements of enzyme activity. From the representations available in the KEGG pathway database [10], we assembled the description of the pathway shown in Fig. 1A (presented in SBGN notation [31]). From this description, we built a deterministic model of the pathway in which catalyzed metabolic transitions were described with Michaelis-Menten kinetics and the autocatalyzed metabolic transitions were described with mass action kinetics. Metabolites play a role in a range of cellular processes and undergo degradation. To capture this each metabolite was also considered to be consumed in an interaction competing with the main pathway, but at a rate lower than its consumption in the main pathway. The competing reactions were modelled with mass action kinetics (not shown in Fig. 1A).

**Figure 1:**
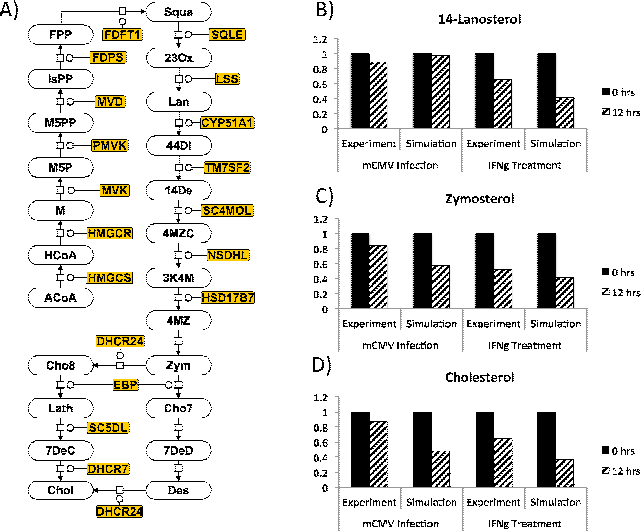
A) The cholesterol pathway represented in SBGN notation, starting with the metabolites Acetyl-Coenzyme A (ACoA) and ending in cholesterol synthesis. B)-D) The normalized concentrations of 14-Lanosterol (B), Zymosterol (C) and Cholesterol (D) at 0 hours (solid, black) and 12 hours (diagonal stripes) after mCMV infection and after IFN*γ* treatment. We show results from experiment and simulation. Experimental measurements were normalized against measurements from a mock time course and simulated measurements were normalized against the concentration at 0hrs.

Parameter values for the cholesterol biosynthesis pathway were taken from the Brenda enzyme database [32]. Where parameters were not known, they were approximated with the mean of the corresponding known parameters. The mean values were 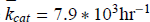 and 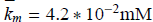. Normalized enzyme levels were inferred from the microarray time courses described above. To obtain an absolute scale for these measurements, we assumed an average number of 5000 enzyme proteins [33] in a region of the cell (the *endoplasmic reticulum*) of volume 10^−141^ [34, 35, 36]. This gave a concentration scale of 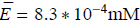 which we took to correspond to the mean normalized enzyme level measurement across both experiments at the start of each time course (mean normalized measurement = 1279.2). We subsequently transformed the normalized expression levels to concentrations using this equivalence and took expression levels to be commensurate with protein levels following the validation described above.

From Supplementary section 2, we can see that, in the limit of low substrate, a Michaelis-Menten interaction acts much like a mass action interaction with a rate constant *k* = *k*_*cat*_ *E*/*k*_*m*_, where *E* is the enzyme concentration. Using 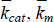 and 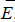, this gave *k* = 156hr^−1^ and we used this value as the rate constant for all the autocatalyzed, mass-action interactions in the main pathway.

In order for the behaviour of the cholesterol biosynthesis pathway activity to dominate over the other dynamical effects, the competing interactions were taken to have mass action rate constants two orders of magnitude smaller than the corresponding mass action constants formed from the main pathway parameters. Thus, for substrate *i*, consumed in a Michaelis-Menten interaction, the corresponding mass action constant would be 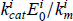 and we assigned to the mass action constant of the competing interaction the value 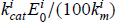, where 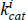 and 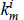 are the turnover and Michaelis-Menten constants of the Michaelis-Menten pathway interaction consuming the metabolite and 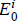 is the enzyme concentration at the start of the time course. Cholesterol was assumed to be consumed at the same rate that it was created in order to avoid accumulation.

A complete list of parameter values is shown in Supplementary sections 3 and 4 and a complete list of normalized enzyme measurements is shown in Supplementary sections 5 and 6.

## 2.3 Modelling strategy

Near saturation, small fluctuations in enzyme concentration lead to small changes of *V*_*max*_ in Michaelis-Menten interactions (*V*_*max*_ = *k*_*cat*_ × enzyme concentration) and this, in turn, leads to large changes in metabolite concentration. Hence, we would expect the pathway to operate in a regime away from saturation where the dynamics would be more stable and robust. Here, we assumed that the pathway was operating away from saturation of the Michaelis-Menten interactions.

We assumed a level of flux into the pathway ∼ 2/3rds of the lowest *V*_*max*_ value obtained across both time courses. From the parameters and the enzyme concentrations at the start of each time course, we calculated the concentrations of each metabolite that would allow the pathway to continue in dynamic equilibrium if no changes were observed in the enzyme concentrations (see Supplementary section 7). The pathway was then numerically integrated, allowing the enzyme concentrations to change in accordance with the concentrations measured in the time course. In the interval between time points, enzyme concentrations were calculated through linear interpolation.

In each interaction of the pathway, the flux was defined and calculated as the rate at which each metabolite is produced. Because this pathway is stoichiometrically trivial, this allowed a direct comparision between all interactions in the pathway. Numerical integration of the pathway was conducted in a two step process, with all the fluxes 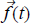 being calculated from metabolite concentrations and then all the metabolite concentrations being updated from the net fluxes. For a general interaction in the interior of a sequential pathway the update rule took the form *m*_*i*_(*t* + Δ*t*) = *m*_*i*_(*t*) + (*f*_*i*-1_(*t*) – *f*_*i*_(*t*))Δ*t*, where *m*_*i*_ is the concentration of metabolite *i*. Modifications to this rule were required at branch points in the pathway and at the start and end. In order to determine a size for Δ*t*, simulations were run in iterations, with values for Δ*t* progressively decreasing. Iterations continued until a value for Δ*t* was reaching at which the results stabilized with no variation in the first four significant figures of the pathway output. This determined the Δ*t* size.

## 3 Results

## 3.1 Model validation

Our first step was to assess the quality of the model by testing whether it behaved in a manner consistent with the observed underlying biology. We did this by comparing the concentrations of three metabolites that cover the pathway at 12 hours following immune challenge.

In Figs. 1B-D, we see normalized metabolite concentrations at 0 and 12 hours following mCMV infection or following IFN*γ* treatment, from both experiment and simulation. Experimentally determined concentrations were normalized against the mock treatment time course; computationally determined concentrations were normalized against the concentration calculated at the start of the time course, determined as part of the initial conditions (see section 2.2).

From a comparison of the experimental and the computationally determined values, we can see that the behaviour of the model is in qualitative agreement with the experimentally observed response of the macrophage cells to mCMV infection and IFN*γ* treatment. This agreement becomes even clearer if we consider the values at 0, 12 and 24 hours post infection or post treatment (Supplementary Fig. 1). This gives us confidence that the model can be used to address questions surrounding the relationship between changes in enzyme levels, metabolite concentrations and flux.

## 3.2 In response to IFN*γ* treatment, the flux through the cholesterol biosynthesis pathway is significantly suppressed in a graduated manner

It is not known how coordinate enzyme control impacts upon flux through the pathway and cholesterol biosynthesis. Thus, we first chose to assess the response of the pathway to the enzyme time courses measured following IFN*γ* treatment.

We simulated the pathway activity by taking the measurements from the microarray time course of macrophage cells following IFN*γ* treatment to represent enzyme concentrations (we have previously shown good correlation between mRNA concentrations and enzyme levels [11]). Fig. 2A shows the resulting profile of flux along the pathway and how this profile develops over the 12 hours. For presentational purposes, we numbered the interactions with 1 representing the input flux and 17 representing cholesterol synthesis (for the full numbering, see Supplementary section 12). The pathway forks at Zymosterol with flux split down both forks. Cholesterol synthesis occurs on both forks and for presentational simplicity, we omitted the details of each fork, but retained the cholesterol synthesis rate, calculated as the sum of the synthesis rates from each fork.

**Figure 2:**
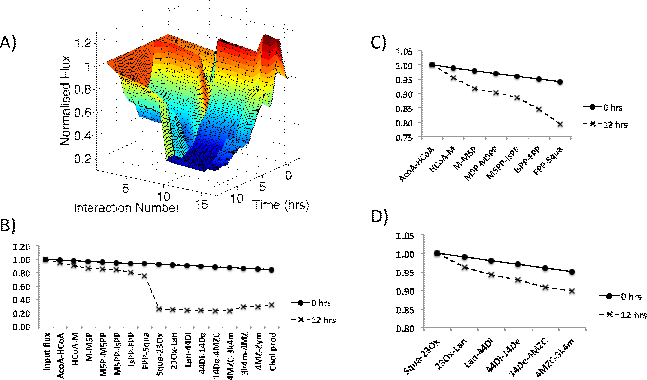
The flux through the cholesterol biosynthesis pathway following treatment with IFN*γ*. A) The development of the flux through the pathway in simulation is shown from 0 hrs to 12 hrs following treatment. Interactions are numbered from 1 (the input flux) to 17 (cholesterol production). For the full numbering, see Supplementary section 12. At 0 hrs, the flux through the pathway is relatively constant. However, by 12 hrs the flux has been significantly suppressed along the pathway. B) The profile of flux through the pathway at 0 hrs and 12 hrs following treatment. These profiles represent cross sections of the surface shown in A). The flux is dramatically reduced in the first 12 hrs. Interactions can be classified as dominant (Squa-23Ox) and non-dominant (the remainder) depending on their degree of impact on the pathway flux. C) The flux through the non-dominant interactions between ACoA-HCoA and FPP-Squa, normalized against the flux through the ACoA-HCoA interaction. The flux through these non-dominant interactions is suppressed more modestly than in the dominant interactions. D) The flux through the non-dominant interactions between Squa-23Ox and 4MZC-3K4M normalized against the flux through Squa-23Ox. The flux through these non-dominant interactions is suppressed more modestly than in the dominant interactions.

From Fig. 2A, we can see that the profile of flux is relatively constant across the pathway at the point of treatment, but that, as time advances, the flux along the pathway becomes suppressed, leading to a much reduced rate of cholesterol synthesis at 12 hours post treatment.

In order to explore the down-regulation in pathway activity further, we examined the cross sections taken at 0 and 12 hours post treatment. The resulting profiles are shown in Fig. 2B. At the point of treatment (0 hrs), we can see that the pathway undergoes a very modest reduction in flux along its length, attributable to the flux lost through the interactions competing for each metabolite. However, as a result of the coordinate down regulation of enzyme activity across the time course, this modest reduction is dramatically amplified.

This profile of flux reduction along the pathway can be considered in terms of (a) dominant interactions, in which the flux reduces significantly and (b) non-dominant interactions, in which the flux reduces more modestly. From Fig. 2B, we can see that the dominant interaction is Squalene-2,3Oxidosqualene (henceforth Squa-23Ox). In order to further explore the degree of suppression in pathway flux that takes place in the non-dominant interactions, we investigated the profile of flux leading up to Squa synthesis and from Squa to 3-keto-4-methyl-zymosterol (henceforth 3k4m), normalizing the flux values against the flux through the first interaction in the sequence. The flux profiles are shown in Figs. 2C and 2D, respectively. In both Figs. 2C and 2D, we can see that the flux through the sequence of interactions is significantly suppressed after 12 hours.

## 3.3 In response to mCMV infection, the flux through the cholesterol biosynthesis pathway shows some suppression in a graduated manner

Ligand activation of the IFN*γ* gamma receptor by the IFN*γ* cytokine is involved in immune activation of macrophages. To compare modulation of the pathway in response to IFN*γ* with its modulation in response to infection, we next modelled the pathway activity using the time course recorded in response to mCMV infection. Fig. 3A shows the resulting profile of flux along the pathway and how this profile developed over time. As in Fig. 2, we numbered the interactions with 1 representing the input flux and 17 representing cholesterol synthesis (see Supplementary section 12). Because the pathway forks at Zymosterol, we omitted the details of the flux along each fork, but retained the cholesterol synthesis rate, calculated as the sum of the synthesis rates on each fork.

**Figure 3:**
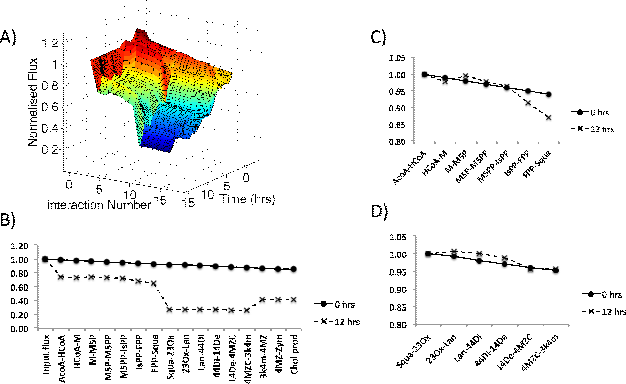
The flux through the cholesterol biosynthesis pathway following infection with mCMV. A) The development of the flux through the pathway in simulation is shown from 0 hrs to 12 hrs post infection. Interactions are numbered from 1 (the input flux) to 17 (cholesterol production). For the full numbering, see Supplementary section 12. At 0 hrs, the flux through the pathway is relatively constant. However, by 12 hrs the flux has been significantly suppressed along the pathway. B) The profile of flux through the pathway at 0 hrs and 12 hrs post infection. These profiles represent cross sections of the surface shown in A). The flux is dramatically reduced in the first 12 hrs following infection. Interactions can be classified as dominant (ACoA-HCoA and Squa-23Ox) and non-dominant (the remainder) depending on their degree of impact on the pathway flux. C) The flux through the non-dominant interactions between ACoA-HCoA and FPP-Squa, normalized against the flux through the ACoA-HCoA interaction. The flux through these non-dominant interactions shows a mild suppression towards the end of the sequence. D) The flux through the non-dominant interactions between Squa-23Ox and 4MZC-3K4M normalized against the flux through Squa-23Ox. The flux through these non-dominant interactions shows no suppression between 0 and 12 hours.

The profile of flux in Fig. 3A is relatively constant across the pathway at the moment of infection, but as time increases, the flux along the pathway reduces, leading to a suppressed pathway and a much reduced rate of cholesterol synthesis. To analyze this response further, we again took cross sections from this surface at 0 and 12 hours post infection. In Fig. 3B, we can see that there is a modest reduction in flux at the point of treatment (0 hours) and that this is amplified as a result of the response to treatment.

As mentioned previously, the profile of flux reduction along the pathway can be considered in terms of interactions that make a dominant contribution to flux reduction and those that make a non-dominant contribution. Interestingly, we see that the distribution of dominant interactions is distinct from the distribution seen in the response to IFN*γ* treatment. Here, the dominant interactions are ACoA-HCoA and Squa-23Ox. In order to explore further the regulation of flux through the non-dominant interactions, we examined the flux profiles between dominant interactions, normalizing the flux through each sequence against the flux through the first interaction in the sequence. The results can be seen in Figs. 3C and D. In the first of these profiles, it is clear that the flux at 12 hours post infection is similar to that at infection, but that some modest reduction does occur towards the end of the sequence as a result of the enzyme regulation. In the second profile, the flux profile clearly does not alter significantly between 0 hours and 12 hours post infection.

### 3.4 Under statin-like intervention, the pathway acquires a step reduction in flux

From Figs. 2 and 3, we can see that the pathway has the ability to respond to perturbation throughout its length. We chose to compare these flux profiles to that which is likely to be obtained when statin-like inhibitors that target the HMGCR enzyme are introduced into the pathway.

Fig. 4A shows the profile of flux obtained in the unperturbed pathway and the profiles at 12 hours following IFN*γ* treatment and mCMV infection. Fig. 4A also shows the profile of flux obtained when the statin-like inhibitor is introduced with a concentration sufficient to suppress cholesterol production to the mean of the production rates for IFN*γ* treatment and mCMV infection at 12 hours. Here we can see that the profile takes a very different form, impacting dramatically upon the interactions upstream of Squa. The profile corresponding to statin-like inhibition is less graduated than in the physiological responses both in terms of the dominant interactions and the non-dominant interactions, and such a flat profile arises because the single HMGCR inhibited interaction takes the entire regulatory burden of reducing the flux along the pathway with the other interactions contributing no more flux reduction than in the unperturbed case.

**Figure 4:**
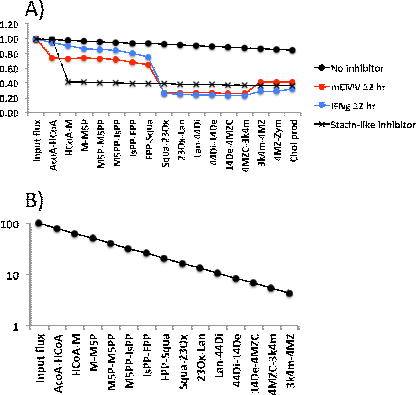
A) The effect of a statin-like inhibitor on pathway activity. The profile of flux at 0 hrs and 12 hrs following mCMV infection and IFN*γ* treatment together with the profile of flux in an unperturbed cell following the introduction of a statin-like inhibitor which targets the enzyme HMGCR. The effect of a statin-like inhibitor is to step down the flux through the interactions catalyzed by HMGCR. This impacts upon the pathway significantly upstream of the point of cholesterol synthesis and creates a flux profile dramatically different to that which arises from the biological response to mCMV infection or IFN*γ* treatment. B) The profile of flux achieved along the pathway when inhibitors concentrations are chosen so that each interaction contributes equally to the regulation of flux (Inhibitor levels listed in Supplementary section 9).

## 4 Discussion

Figs. 2, 3 and 4 give us insight into how the cholesterol biosynthesis pathway operates and its various modes of regulation. Fig. 2 shows that, in response to IFN*γ* treatment, the pathway undergoes a reduction in flux that is mediated by both the dominant and non-dominant interactions. The distribution of flux at the point of IFN*γ* treatment shows a very mild reduction along the pathway. This is attributable to the flux lost through the off-pathway, competing interactions. However, at 12 hours post treatment, we can see that the flux becomes significantly suppressed along the pathway due to the coordinate enzyme reduction. This occurs because a reduction in the enzyme concentration in an interaction has the effect of decreasing the rate of the interaction. This, in turn, increases the proportion of flux shunted through the off-pathway interaction competing for the same metabolite. At 12 hours post treatment, we can see that most of the interactions on the pathway play a suppressive role. This implies that, for most of the metabolites, there is an increase in the proportion of flux shunted through the off-pathway interactions, if flux conservation is to be maintained.

In Fig. 3, we can see that the pathway also undergoes a reduction in flux in response to mCMV infection. In this case, the reduction is mediated mostly by the dominant interactions with very few non-dominant interactions contributing to the suppression of flux between 0 and 12 hours post infection. Here, we infer that the coordinate enzyme control leads to an increased proportion of flux being shunted through the off-pathway interactions that compete for metabolites with the dominant interactions. However, we see little suppression of flux through the non-dominant interactions between 0 and 12 hours post infection, implying that the proportion of flux being shunted through the off-pathway interactions, competing for the same metabolites, stays approximately the same as at the point of infection.

In Fig. 4A, we see that a statin-like pharmaceutical intervention comparable to the immune-led responses introduces a profile of flux with a significantly different shape to that of the immune led response. A statin-like intervention leads to a step down in flux towards the start of the pathway with the result that the flux passing through many of the interactions in the upper half of the pathway is significantly lower than the levels observed in immune-led responses. We can infer that a statin-like intervention shunts a significantly increased proportion of flux through the off-pathway interaction competing for HCoA. In the remaining interactions, no significant change in the proportion of flux passing through the main pathway interactions occurs.

If we consider the flux profiles shown in Fig. 4A, we see that the introduction of a statin-like inhibitor steps down the flux at the interaction HCoA-M and that this has consequences for many of the interactions downstream of HCoA-M. In the upper regions of the pathway, the level of flux after the introduction of a statin-like inhibitor is well below that achieved in immune-led responses. This is particularly significant for the interactions that consume and produce Isopentyl-PP (IsPP) and Farnesyl-PP (FPP) as these metabolites are also consumed in the prenylation and dolichylation non-sterol arms that fork from the main sterol path-way (***cite Mazein et al, same issue***)[**?**, 19, 20]. Hence, the difference in profiles between statin-like intervention and the immune-led regulation is likely to have significant consequences for these non-sterol arms due to the difference in flux passing through the branch point in the pathway. Off target effects are known to be a particular concern for statin based treatments [30, 29] and the difference in flux profiles may provide a clue as to why this is the case.

If we consider the profiles of flux associated with coordinate enzyme regulation and with single enzyme regulation, it is clear that the coordinate regulation provides a more robust strategy for regulating the pathway. Single enzyme regulation offers no protection against metabolite surges occurring downstream of the regulated interaction. Therefore, distributing flux reduction over multiple interactions, rather than a single interaction, protects the regulatory mechanism from cellular perturbations. The regulation of the cholesterol biosynthesis pathway by IFN*γ* has been shown to be part of an antiviral response and a distributed flux reduction also increases the difficulty for a pathogen to manipulate the pathway to proviral effect. For these reasons, it is likely that coordinate control has enjoyed a selective advantage in the evolution of immune regulation.

From Figs. 2 and 3, we can see that all the interactions have the potential to play a regulatory role. Therefore, an important question to ask is whether a combination of inhibitors can be chosen such that the flux reduction is distributed evenly throughout the pathway. Such a regulatory structure would be optimally robust. This could be achieved through control at the transcriptional level, tuning SREBP2 activity, or at the enzyme level using pharmaceutical inhibition or miRNA induced degradation. Such an arrangement can be achieved in the models described above by using the combination of inhibitors with levels described in Supplementary section 9. The resulting flux profile is shown in Fig. 4B. Here, the inhibitor concentrations have been chosen so that, for each metabolite, the proportion of flux shunted down the off-pathway interaction to flux continuing down the main pathway is the exactly same. In addition to the superior robustness of such a distributed regulation, this profile of flux would also show greater specificity than a stepped, statin-like treatment as the upstream interactions would be significantly less flux suppressed, reducing the impact on off-pathway interactions. It is possible to speculate that a regulatory arrangement that targets a single interaction towards the end of the pathway would demonstrate even greater specificity to cholesterol as it would create a flux profile in which the flux remains at unregulated levels for most of the pathway, but steps down immediately before the point of cholesterol synthesis. However, such a regulatory structure would not enjoy the robustness of multiple interaction regulation. The improved specificity and robustness that comes with multiple interaction regulation suggests that future pharmacological therapies may be able to act more efficiently and with fewer side effects, if they are designed either to mimic the physiological response of IFN*γ* treatment or to create a distributed regulation of flux comparable to Fig. 4B.

Further interesting detail can be observed when we allow our simulations to run up to 24 hours post infection or post IFN*γ* treatment. At 24 hours we had access to only a limited number of QPCR measurements for a subset of the enzymes on the pathway, so to provide values for the unmeasured enzymes at 24 hours, we assumed that the concentration did not change from the 12 hour time point. From Supplementary Fig. 1, we can see that qualitative agreement between our simulated metabolite levels and the measured values improves in the 12-24 hour window, further supporting the use of this model to explore the dynamics of the cholesterol biosynthesis pathway. Exploring the flux profiles at 24 hours post IFN*γ* treatment and post mCMV infection (Supplementary Figs. 2 and 3, respectively), we can see that there is a difference in the relative timing of the flux suppression. In mCMV infection, pathway flux is suppressed at 12 hours post infection and continues to be further suppressed over the 12 to 24 hour interval. However, in response to IFN*γ* treatment, we see that the pathway starts to recover its pretreated profile of flux in the 12 to 24 hour window. The simplest explanation for this difference would be that there is an inherent delay between mCMV infection and the induction of IFN*γ* signalling within cell populations. This would predict that running the mCMV infection experiments for a longer duration would yield enzyme expression measurements that would lead to a rise in pathway flux similar to that observed in IFN*γ* treatment. Although at 12 hours post mCMV infection there was little evidence of the non-dominant interactions contributing to pathway regulation, it is also worth noting that, at 24 hours post mCMV infection, the non-dominant interactions start to show a modest suppression of flux comparable to that seen in the 0-12 hour window following IFN*γ* treatment. This further supports the idea that the two profiles are related.

The flux profiles shown in Figs. 2A and 3A and in Supplementary Figs. 2A and 3A exhibit interesting features. The broad trend is towards pathway suppression. However, there are points at which the flux exceeds unperturbed levels and where the flux downstream exceeds the flux into the pathway. This may be attributable, at least in part, to technical noise in the time course microarray measurements. However, we cannot rule out that this is biological. Points where the flux consuming a metabolite exceeds the flux producing it correspond to a reduction in the concentration of a metabolite (data not shown). In Figs 2B and 3B, we see that there is a rise in flux at the 3K4M-4MZ interaction. In our simulations, this corresponded to a reduction in the concentration of 3K4M (data not shown).

Making quantitative predictions of pathway function requires a comprehensive set of high confidence parameters. Without such a parameter set, we are restricted to qualitative observations. Despite the importance of the cholesterol biosynthesis pathway to innate immunity and cardiovascular health and despite its value to industry as the target pathway of the statin class of drugs, we can see from the Table in Supplementary section 3 that the parameterization of this pathway is largely incomplete. This is a surprising result. The lack of a rigorous parameterization impedes studies of pathway behaviour and the development of more advanced scientific and therapeutic interventions. The incomplete nature of the parameterization led us to use estimates for the unknown parameters and this has implications for the flux profiles we obtained in simulations. The particular arrangement of dominant interactions and non-dominant interactions, for example, is parameter dependent and we can see from Supplementary section 3 that the two interactions identified as playing a dominant role in flux reduction (ACoA-HCoA and Squa-23Ox) are those with the lowest turnover (*k*_*cat*_) values. Accurate predictions of the precise arrangement of dominant and non-dominant interactions require a complete, high confidence parameter set. However, under the assumptions made in the model, it is a parameter independent result that all the interactions can contribute a modulation of flux.

The coordinate down regulation of enzyme activity has been demonstrated to be an innate, immune, anti-viral response to mCMV infection mediated by IFN*γ*, rather than a pathogenic intervention [11] and so it would appear that flux reduction is also part of the innate immune response. The cholesterol biosynthesis pathway has been demonstrated to be critical to the optimal viral infection of a further range of viruses (Hepatitis C (HCV), HIV, Japanese Encephalitis (JEV), West Nile (WNV), Dengue (DENV), Measles viruses (MV) and African Swine Fever Viruses (ASFV) [6, 7, 11, 12, 13, 14, 15, 18]). Since the flux suppression is in response to IFN*γ*, a general immunological cytokine, it is possible that such a flux suppression might be part of a general antiviral response.

## 5 Conclusion

Here, we have shown, with a simple dynamical model, that the coordinate down regulation of the enzymes in the cholesterol biosynthesis pathway in macrophages, observed in response to mCMV infection and IFN*γ*, treatment leads to a graduated reduction in flux along the pathway that can be considered in terms of dominant interactions that significantly suppress the flux and non-dominant interactions that make modest, but not insignificant contributions to flux suppression. The model was built using a combination of Michaelis-Menten and mass action interactions and was validated using experimental measurements of metabolite concentrations at a range of time points. Our result suggests that all the interactions in the pathway could be candidate intervention points for future therapeutic strategies.

We were able to compare the impact on flux of coordinate regulation which acts to regulate multiple interactions to the impact of statin-like inhibitors which regulate a single enzyme. This allowed us to explore the advantages of multiple interaction regulation over single enzyme regulation. The graduated profile generated by coordinate regulation enjoys greater robustness and greater specificity and we highlight the importance of this for the prenylation and dolichylation non-sterol arms that fork from the cholesterol biosynthesis pathway. We suggest that such a multiple interaction regulation could be exploited in future pharmaceutical therapies in order to improve their efficacy and specificity, perhaps mimicking the regulation of the pathway observed in immunological responses.

## 6 Acknowledgments

This work was supported by the Wellcome Trust (WT066784) programme grant and BBSRC to PG. The Centre for Systems Biology at Edinburgh is a centre for Integrative Systems Biology supported by the BBSRC and EPSRC (BB/D019621/1).

## 7 Author Contribution

AM, SW, HG, KAR and PG compiled a description of the cholesterol biosynthesis pathway. SW, MLG, JH and PG modelled the pathway. SW, LL and MLG compiled parameters for the pathway. MB, GS and MRW completed the experiments. Analysis was completed by SW and PG. SW, KAR and PG wrote the manuscript with feedback from all authors.

## Supplementary Material - A model of flux regulation in the cholesterol biosynthesis pathway: Immune mediated graduated flux reduction versus statin-like led stepped flux reduction

### 1 Abbreviations used in Fig. 1A

1 Abbreviations used in Fig. 1A.
| Acronym | Metabolite |
| --- | --- |
| ACoA | Acetyl-CoA |
| HCoA | HMG-CoA |
| M | Mevalonate |
| M5P | Mevalonate-5P |
| M5PP | Mevalonate-5PP |
| IsPP | Isopentyl-PP |
| FPP | Farnesyl-PP |
| Squa | Squalene |
| 23Ox | 2,3 oxydosqualene |
| Lan | Lanosterol |
| 44Di | 4,4 dimethyl-cholesta-8,14,24-trienol |
| 14De | 14-demethyl-lanosterol |
| 4MZC | 4-methylzymosterol-carboxylate |
| 3K4M | 3-keto-4-methyl-zymosterol |
| 4MZ | 4-methylzymosterol |
| Zym | Zymosterol |
| Cho7 | Cholesta-7,24-dien-3beta-ol |
| 7DeD | 7-dehydro-desmosterol |
| Des | Desmosterol |
| Cho8 | Cholesta-8,en-3beta-ol |
| Lath | Lathosterol |
| 7DeC | 7 dehydro-cholesterol |
| Chol | Cholesterol |

### 2. Relating Michaelis-Menten interactions to Mass-action interactions

For a Michaelis-Menten interaction of the form

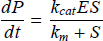

where *k*_*cat*_ is the turnover parameter, *k*_*m*_ is the Michaelis-Menten constant, *E* is the enzyme concentration and *S* is the substrate concentration, we want to investigate the consequences of *S* being small (ie much less that *k*_*m*_). If we define a small value for *S* as Δ*S* (where Δ*S* << *k*_*m*_), we have

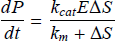

Dividing the numerator and denominator by *k*_*m*_ gives

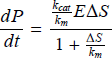

We have 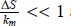 and so we can use the identity that

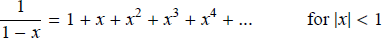

Thus

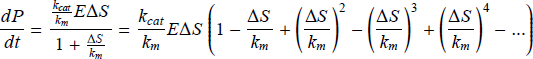

To second order in Δ*S* , this is

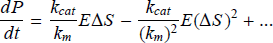

As 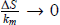 all but the first order term make negligible contributions, meaning that

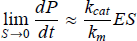

### 3 Parameters gathered from the Brenda enzyme database and used in our simulations

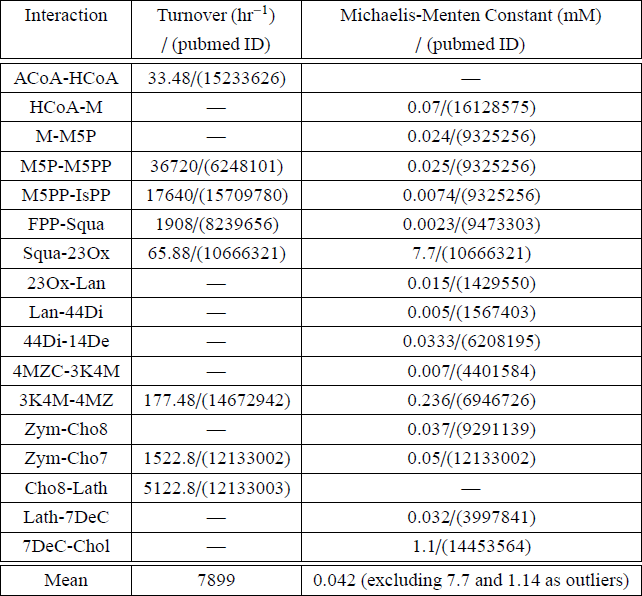

Retrieved 27th October, 2009.

### 4 Estimated parameters

| Interaction | Turnover ($\text{hr}^{-1}$)<br>/ (pubmed ID) | Michaelis-Menten Constant (mM)<br>/ (pubmed ID) |
| --- | --- | --- |
| ACoA-HCoA | — | 0.042 |
| HCoA-M | 7900 | — |
| M-M5P | 7900 | — |
| IsPP-FPP | 7900 | 0.042 |
| 23Ox-Lan | 7900 | — |
| Lan-44Di | 7900 | — |
| 44Di-14De | 7900 | — |
| 14De-4MZC | 7900 | 0.042 |
| 4MZC-3K4M | 7900 | — |
| Des-Chol | 7900 | 0.042 |
| Zym-Cho8 | 7900 | — |
| Cho8-Lath | — | 0.042 |
| Lath-7DeC | 7900 | — |
| 7DeC-Chol | 7900 | — |

| Interaction | Mass action constant ($\text{hr}^{-1}$) |
| --- | --- |
| 4MZ-Zym | 156 |
| Cho7-7DeD | 156 |
| 7DeD-Des | 156 |

| Interaction | Rate constant of competing interaction (hr<sup>-1</sup>) - IFNγ treatment |
| --- | --- |
| ACoA-HCoA | 0.0134 |
| HCoA-M | 0.1977 |
| M-M5P | 0.1858 |
| M5P-M5PP | 8.8154 |
| M5PP-IsPP | 2.0107 |
| IsPP-FPP | 8.6358 |
| FPP-Squa | 15.5664 |
| Squa-23Ox | 0.0004 |
| 23Ox-Lan | 0.3178 |
| Lan-44Di | 1.1174 |
| 44Di-14De | 0.6450 |
| 14De-4MZC | 6.7600 |
| 4MZC-3K4M | 0.5712 |
| 3K4M-4MZ | 0.0011 |
| 4MZ-Zym | 1.5600 |
| Zym-Cho7 | 0.2680 |
| Cho7-7DeD | 1.5600 |
| 7DeD-Des | 1.5600 |
| Des-Chol | 0.1855 |
| Zym-Cho8 | 0.2680 |
| Cho8-Lath | 1.3027 |
| Lath-7DeC | 1.3936 |
| 7DeC-Chol | 0.0296 |

| Interaction | Rate constant of competing interaction ($\text{hr}^{-1}$) - mCMV infection |
| --- | --- |
| ACoA-HCoA | 0.00745 |
| HCoA-M | 0.18893 |
| M-M5P | 0.16232 |
| M5P-M5PP | 8.32940 |
| M5PP-IsPP | 1.71684 |
| IsPP-FPP | 8.57849 |
| FPP-Squa | 18.43532 |
| Squa-23Ox | 0.00003 |
| 23Ox-Lan | 0.26313 |
| Lan-44Di | 0.77913 |
| 44Di-14De | 0.60956 |
| 14De-4MZC | 5.82151 |
| 4MZC-3K4M | 0.31487 |
| 3K4M-4MZ | 0.00127 |
| 4MZ-Zym | 1.56000 |
| Zym-Cho7 | 0.2008 |
| Cho7-7DeD | 1.56000 |
| 7DeD-Des | 1.56000 |
| Des-Chol | 0.16598 |
| Zym-Cho8 | 0.2008 |
| Cho8-Lath | 0.85392 |
| Lath-7DeC | 0.96270 |
| 7DeC-Chol | 0.02307 |

### 5 Normalized enzyme activity time course following infection with mCMV

5 Normalized enzyme activity time course following infection with mCMV.
| Time (hr) | HMGCR | MVK | PMVK | MVD | FDPS | FDFT1 | SQLE | DHCR7 |
| --- | --- | --- | --- | --- | --- | --- | --- | --- |
| 0 | 258 | 76 | 874 | 111 | 7029 | 3425 | 486 | 495 |
| 0.5 | 253 | 68 | 855 | 106 | 6990 | 3322 | 428 | 436 |
| 1 | 255 | 54 | 790 | 91 | 6568 | 3509 | 490 | 339 |
| 1.5 | 271 | 38 | 735 | 76 | 6036 | 3346 | 515 | 279 |
| 2 | 267 | 35 | 639 | 65 | 5082 | 2942 | 530 | 297 |
| 2.5 | 271 | 39 | 590 | 70 | 4612 | 2411 | 483 | 280 |
| 3 | 259 | 47 | 541 | 66 | 4186 | 1861 | 462 | 259 |
| 3.5 | 287 | 44 | 524 | 64 | 4387 | 1674 | 466 | 175 |
| 4 | 293 | 50 | 515 | 68 | 4666 | 1445 | 396 | 215 |
| 4.5 | 314 | 51 | 501 | 74 | 5062 | 1341 | 375 | 231 |
| 5 | 304 | 66 | 522 | 80 | 4848 | 1154 | 304 | 296 |
| 5.5 | 304 | 64 | 535 | 71 | 4404 | 1022 | 309 | 322 |
| 6 | 290 | 77 | 550 | 73 | 3468 | 997 | 282 | 334 |
| 6.5 | 307 | 63 | 574 | 65 | 3335 | 1029 | 318 | 314 |
| 7 | 318 | 73 | 587 | 68 | 3224 | 1114 | 310 | 306 |
| 7.5 | 317 | 58 | 613 | 61 | 3573 | 940 | 281 | 282 |
| 8 | 300 | 62 | 580 | 67 | 3303 | 919 | 271 | 304 |
| 8.5 | 289 | 55 | 562 | 64 | 3070 | 825 | 260 | 266 |
| 9 | 286 | 66 | 580 | 65 | 2713 | 927 | 217 | 244 |
| 9.5 | 273 | 56 | 546 | 56 | 2516 | 895 | 207 | 219 |
| 10 | 254 | 42 | 517 | 45 | 2434 | 872 | 205 | 208 |
| 10.5 | 232 | 31 | 464 | 43 | 2233 | 840 | 243 | 215 |
| 11 | 212 | 39 | 446 | 48 | 1915 | 764 | 187 | 205 |
| 11.5 | 199 | 50 | 445 | 55 | 1516 | 731 | 154 | 193 |
| 12 | 196 | 66 | 459 | 67 | 1382 | 658 | 141 | 187 |
| 24 | 105 | 66 | 459 | 44 | 1639 | 1087 | 105 | 631 |

| Time (hr) | HMGCS | LSS | CYP51A1 | TM7SF2 | SC4MOL | NSDHL | HSD17B7 | DHCR24 |
| --- | --- | --- | --- | --- | --- | --- | --- | --- |
| 0 | 1441 | 77 | 76 | 396 | 4770 | 43 | 261 | 136 |
| 0.5 | 1510 | 72 | 74 | 402 | 4570 | 44 | 253 | 117 |
| 1 | 1707 | 85 | 72 | 391 | 4931 | 52 | 254 | 115 |
| 1.5 | 1809 | 92 | 69 | 373 | 5256 | 63 | 258 | 108 |
| 2 | 1649 | 98 | 60 | 312 | 5158 | 68 | 242 | 102 |
| 2.5 | 1389 | 90 | 61 | 299 | 4859 | 72 | 250 | 97 |
| 3 | 1307 | 88 | 61 | 287 | 3901 | 79 | 238 | 91 |
| 3.5 | 1502 | 85 | 67 | 323 | 3688 | 77 | 255 | 89 |
| 4 | 1538 | 87 | 61 | 296 | 3320 | 59 | 216 | 84 |
| 4.5 | 1665 | 81 | 62 | 322 | 3169 | 52 | 208 | 84 |
| 5 | 1309 | 79 | 65 | 319 | 2887 | 58 | 173 | 89 |
| 5.5 | 1101 | 74 | 67 | 344 | 2359 | 68 | 169 | 90 |
| 6 | 853 | 71 | 65 | 314 | 2056 | 65 | 155 | 89 |
| 6.5 | 877 | 74 | 64 | 330 | 1875 | 64 | 169 | 84 |
| 7 | 928 | 74 | 66 | 341 | 1814 | 67 | 173 | 90 |
| 7.5 | 893 | 72 | 63 | 347 | 1832 | 63 | 173 | 90 |
| 8 | 799 | 68 | 55 | 319 | 1694 | 59 | 163 | 84 |
| 8.5 | 678 | 60 | 51 | 307 | 1615 | 56 | 160 | 77 |
| 9 | 628 | 57 | 50 | 293 | 1603 | 58 | 149 | 74 |
| 9.5 | 672 | 56 | 54 | 276 | 1689 | 63 | 151 | 79 |
| 10 | 710 | 59 | 52 | 260 | 1764 | 65 | 139 | 81 |
| 10.5 | 724 | 56 | 53 | 247 | 1659 | 66 | 150 | 76 |
| 11 | 605 | 53 | 53 | 248 | 1460 | 61 | 140 | 73 |
| 11.5 | 558 | 47 | 55 | 227 | 1428 | 56 | 139 | 63 |
| 12 | 574 | 52 | 63 | 225 | 1426 | 59 | 133 | 69 |
| 24 | 301 | 44 | 22 | 225 | 1334 | 21 | 149 | 140 |

| Time (hr) | EBP | SC5DL |
| --- | --- | --- |
| 0 | 1079 | 601 |
| 0.5 | 1533 | 583 |
| 1 | 1505 | 664 |
| 1.5 | 1380 | 771 |
| 2 | 1258 | 809 |
| 2.5 | 1192 | 792 |
| 3 | 1277 | 693 |
| 3.5 | 1402 | 757 |
| 4 | 1402 | 734 |
| 4.5 | 1359 | 739 |
| 5 | 1182 | 623 |
| 5.5 | 1244 | 525 |
| 6 | 1272 | 452 |
| 6.5 | 1497 | 459 |
| 7 | 1628 | 500 |
| 7.5 | 1709 | 471 |
| 8 | 1778 | 465 |
| 8.5 | 1691 | 413 |
| 9 | 1655 | 435 |
| 9.5 | 1613 | 431 |
| 10 | 1647 | 463 |
| 10.5 | 1659 | 454 |
| 11 | 1493 | 417 |
| 11.5 | 1298 | 372 |
| 12 | 1224 | 359 |
| 24 | 1224 | 359 |

### 6 Normalized enzyme activity time course following treatment with IFN*γ*

6 Normalized enzyme activity time course following treatment with IFN $\gamma$ .
| Time (hr) | HMGCR | MVK | PMVK | MVD | FDPS | FDFT1 | SQLE | DHCR7 |
| --- | --- | --- | --- | --- | --- | --- | --- | --- |
| 0 | 270 | 87 | 925 | 130 | 7076 | 2892 | 677 | 635 |
| 0.5 | 296 | 75 | 902 | 134 | 6391 | 3508 | 647 | 577 |
| 1 | 277 | 83 | 886 | 149 | 7127 | 3302 | 768 | 680 |
| 1.5 | 320 | 84 | 1016 | 137 | 7821 | 3668 | 810 | 661 |
| 2 | 288 | 94 | 1014 | 140 | 7617 | 2795 | 811 | 753 |
| 2.5 | 286 | 74 | 932 | 109 | 5545 | 2561 | 648 | 568 |
| 3 | 245 | 64 | 821 | 96 | 4392 | 1956 | 539 | 537 |
| 3.5 | 247 | 55 | 872 | 69 | 4355 | 1934 | 464 | 423 |
| 4 | 271 | 66 | 1107 | 69 | 5263 | 1834 | 407 | 409 |
| 4.5 | 254 | 69 | 1193 | 72 | 5082 | 1678 | 375 | 372 |
| 5 | 258 | 87 | 1347 | 89 | 5306 | 1579 | 344 | 360 |
| 5.5 | 247 | 78 | 1299 | 87 | 4728 | 1419 | 326 | 323 |
| 6 | 241 | 85 | 1301 | 90 | 5479 | 1175 | 271 | 311 |
| 6.5 | 228 | 66 | 1190 | 81 | 5239 | 928 | 331 | 252 |
| 7 | 211 | 63 | 1169 | 81 | 5324 | 807 | 203 | 259 |
| 7.5 | 207 | 55 | 1110 | 75 | 4373 | 745 | 197 | 212 |
| 8 | 219 | 67 | 1003 | 88 | 3942 | 756 | 188 | 206 |
| 8.5 | 224 | 80 | 957 | 84 | 3840 | 694 | 172 | 168 |
| 9 | 219 | 80 | 834 | 79 | 3189 | 600 | 152 | 157 |
| 9.5 | 190 | 72 | 857 | 74 | 2811 | 515 | 159 | 162 |
| 10 | 174 | 57 | 789 | 64 | 2264 | 465 | 167 | 166 |
| 10.5 | 186 | 70 | 778 | 71 | 2227 | 527 | 200 | 178 |
| 11 | 186 | 69 | 679 | 60 | 1993 | 521 | 205 | 173 |
| 11.5 | 178 | 67 | 609 | 61 | 1833 | 496 | 202 | 182 |
| 12 | 156 | 58 | 602 | 62 | 1544 | 450 | 185 | 180 |
| 24 | 99 | 58 | 602 | 84 | 3870 | 1667 | 345 | 631 |

| Time (hr) | HMGCS | LSS | CYP51A1 | TM7SF2 | SC4MOL | NSDHL | HSD17B7 | DHCR24 |
| --- | --- | --- | --- | --- | --- | --- | --- | --- |
| 0 | 2594 | 93 | 109 | 419 | 5539 | 78 | 227 | 152 |
| 0.5 | 2491 | 104 | 117 | 410 | 5657 | 89 | 242 | 131 |
| 1 | 2527 | 104 | 120 | 414 | 5593 | 74 | 270 | 138 |
| 1.5 | 1930 | 99 | 151 | 497 | 6129 | 70 | 324 | 122 |
| 2 | 1740 | 92 | 150 | 494 | 5924 | 63 | 326 | 123 |
| 2.5 | 1364 | 85 | 129 | 422 | 5303 | 68 | 247 | 105 |
| 3 | 1663 | 75 | 97 | 334 | 4205 | 59 | 201 | 89 |
| 3.5 | 1803 | 65 | 94 | 331 | 3805 | 51 | 185 | 77 |
| 4 | 1915 | 66 | 116 | 397 | 3919 | 50 | 224 | 84 |
| 4.5 | 1437 | 65 | 123 | 406 | 3693 | 50 | 226 | 90 |
| 5 | 1131 | 71 | 127 | 435 | 3439 | 61 | 219 | 97 |
| 5.5 | 966 | 65 | 120 | 421 | 3130 | 57 | 200 | 86 |
| 6 | 814 | 71 | 138 | 438 | 2694 | 66 | 186 | 85 |
| 6.5 | 581 | 64 | 145 | 425 | 2324 | 60 | 172 | 87 |
| 7 | 476 | 61 | 153 | 427 | 2028 | 63 | 159 | 92 |
| 7.5 | 500 | 55 | 146 | 420 | 1923 | 59 | 138 | 102 |
| 8 | 542 | 57 | 172 | 448 | 1849 | 60 | 146 | 99 |
| 8.5 | 545 | 57 | 202 | 475 | 1676 | 64 | 146 | 96 |
| 9 | 501 | 58 | 203 | 450 | 1714 | 66 | 153 | 89 |
| 9.5 | 513 | 56 | 175 | 420 | 1726 | 67 | 144 | 83 |
| 10 | 535 | 49 | 145 | 367 | 1811 | 60 | 130 | 77 |
| 10.5 | 607 | 58 | 109 | 358 | 1962 | 68 | 136 | 83 |
| 11 | 643 | 50 | 75 | 332 | 2218 | 63 | 122 | 67 |
| 11.5 | 662 | 50 | 52 | 313 | 2402 | 60 | 118 | 64 |
| 12 | 679 | 41 | 49 | 283 | 2421 | 57 | 101 | 55 |
| 24 | 1221 | 58 | 65 | 283 | 5596 | 80 | 136 | 140 |

| Time (hr) | EBP | SC5DL |
| --- | --- | --- |
| 0 | 1646 | 870 |
| 0.5 | 1696 | 908 |
| 1 | 1815 | 911 |
| 1.5 | 2303 | 1026 |
| 2 | 2444 | 995 |
| 2.5 | 2033 | 899 |
| 3 | 1671 | 731 |
| 3.5 | 1610 | 694 |
| 4 | 1870 | 683 |
| 4.5 | 1944 | 662 |
| 5 | 1924 | 638 |
| 5.5 | 1914 | 657 |
| 6 | 2035 | 606 |
| 6.5 | 2202 | 511 |
| 7 | 2239 | 468 |
| 7.5 | 2233 | 460 |
| 8 | 2020 | 512 |
| 8.5 | 1953 | 499 |
| 9 | 1703 | 454 |
| 9.5 | 1703 | 378 |
| 10 | 1780 | 319 |
| 10.5 | 1820 | 302 |
| 11 | 1902 | 290 |
| 11.5 | 1805 | 274 |
| 12 | 1848 | 243 |
| 24 | 1848 | 243 |

### 7 Calculating initial conditions

The most general case is an interaction in which the substrate is consumed in a Michaelis-Menten interaction and an off-pathway, competing mass action interaction. If *F* is the rate at which a substrate is formed, the pathway will be at dynamic equilibrium if

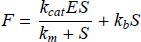

where *k*_*b*_ is the mass action rate constant for the off-pathway interaction. After some rearrangement, this yields the following quadratic equation

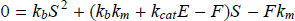

which, from the monotonicity of the mass action term and the Michaelis-Menten terms, can be seen to have one positive solution.

Where an interaction is mass action in form, we have dynamic equilibrium when

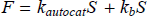

This can be rearranged to

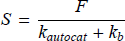

Zymosterol is consumed in two Michaelis-Menten interactions. Here, dynamic equilibrium is established when

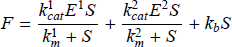

After some rearrangement, this yields the following cubic equation

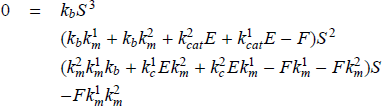

which, from the monotonicity of the two Michaelis-Menten terms and the mass action term, can be seen to have one positive solution.

### 8 Metabolite levels at 0, 12 and 24 hours post infection or post treatment

**Figure 1:**
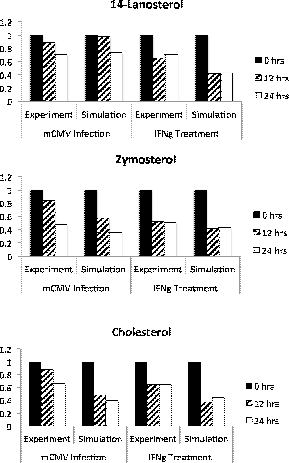
Metabolite levels at 0, 12 and 24 hours post infection or post treatment. The normalized concentrations of 14-Lanosterol, Zymosterol and Cholesterol at 0 hours (solid, black), 12 hours (diagonal stripes) and 24 hours (open boxes) after mCMV infection and after IFN*γ* treatment. We show results from experiment and simulation. Experimental measurements were normalized against measurements from a mock time course and simulated measurements were normalized against the concentration at 0hrs.

### 9 Inhibitor levels responsible for the flux profile in Fig. 4B

| Interaction | Inhibitor to $K_i$ ratio |
| --- | --- |
| ACoA-HCoA | 22.94 |
| HCoA-M | 23.97 |
| M-M5P | 23.91 |
| M5P-M5PP | 24.00 |
| M5PP-IsPP | 23.98 |
| IsPP-FPP | 24.00 |
| FPP-Squa | 24.00 |
| Squa-23Ox | 23.57 |
| 23Ox-Lan | 23.98 |
| Lan-44Di | 23.99 |
| 44Di-14De | 24.00 |
| 14De-4MZC | 24.00 |
| 4MZC-3K4M | 23.99 |
| 3K4M-4MZ | 23.87 |

### 10 Flux profiles from 0 to 24 hours post IFN*γ* treatment

**Figure 2:**
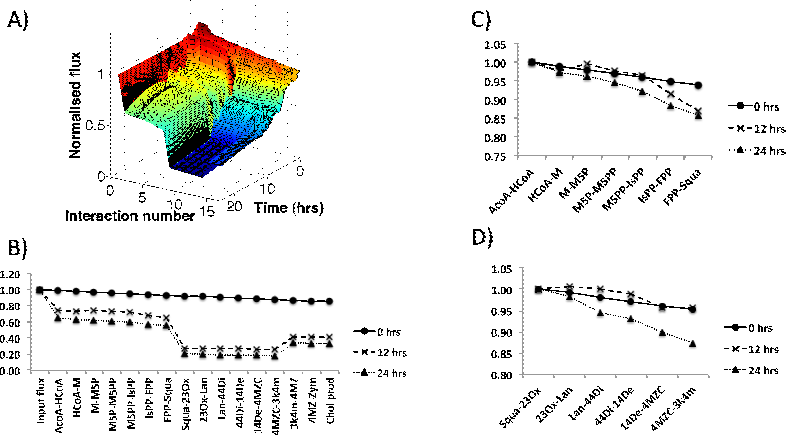
Flux profiles from 0 to 24 hours post IFN*γ* treatment. The flux through the cholesterol biosynthesis pathway following treatment with IFN*γ*. A) The development of the flux through the pathway in simulation is shown from 0 hrs to 24 hrs following treatment. Interactions are numbered from 1 (the input flux) to 17 (cholesterol production). For the full numbering, see Supplementary section 12. At 0 hrs, the flux through the pathway is relatively constant. However, by 12 hrs the flux has been significantly suppressed along the pathway. By 24 hrs, the pathway has started to show a modest recovery. B) The profile of flux through the pathway at 0 hrs, 12 hrs and 24 hrs following treatment. These profiles represent cross sections of the surface shown in A). The flux is dramatically reduced in the first 12 hrs with a modest increase occurring between 12 and 24 hrs following treatment. Interactions can be classified as dominant (Squa-23Ox) and non-dominant (the remainder) depending on their degree of impact on the pathway flux. C) The flux through the non-dominant interactions between ACoA-HCoA and FPP-Squa, normalized against the flux through the ACoA-HCoA interaction. D) The flux through the non-dominant interactions between Squa-23Ox and 4MZC-3K4M normalized against the flux through the interaction Squa-23Ox.

### 11 Flux profiles from 0 to 24 hours post mCMV infection

**Figure 3:**
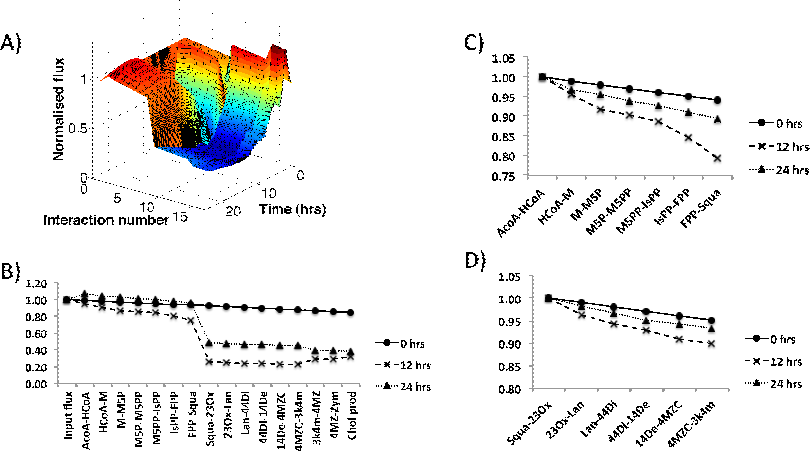
Flux profiles from 0 to 24 hours post mCMV infection. The flux through the cholesterol biosynthesis pathway following infection with mCMV. A) The development of the flux through the pathway in simulation is shown from 0 hrs to 24 hrs post infection. Interactions are numbered from 1 (the input flux) to 17 (cholesterol production). For the full numbering, see Supplementary section 12. At 0 hrs, the flux through the pathway is relatively constant. However, by 24 hrs the flux has been significantly suppressed along the pathway. B) The profile of flux through the pathway at 0 hrs, 12 hrs and 24 hrs post infection. These profiles represent cross sections of the surface shown in A). The flux is dramatically reduced in the first 12 hrs with a further reduction occurring between 12 and 24 hrs following infection. Interactions can be classified as dominant (ACoA-HCoA and Squa-23Ox) and non-dominant (the remainder) depending on their degree of impact on the pathway flux. C) The flux through the non-dominant interactions between ACoA-HCoA and FPP-Squa, normalized against the flux through the ACoA-HCoA interaction. The flux through these non-dominant interactions shows a mild suppression between 0 hrs and 12 hrs, but a more significant suppression between 12 hr sand 24 hrs. D) The flux through the non-dominant interactions between Squa-23Ox and 4MZC-3K4M normalized against the flux through the interaction Squa-23Ox. The flux through these interactions shows no suppression between 0 hrs and 12 hrs, but a significant suppression between 12 hrs and 24 hrs.

### 12 Interaction numbering

|  |  |
| --- | --- |
| 1 | Input flux |
| 2 | AcoA-HCoA |
| 3 | HCoA-M |
| 4 | M-M5P |
| 5 | M5P-M5PP |
| 6 | M5PP-IsPP |
| 7 | IsPP-FPP |
| 8 | FPP-Squa |
| 9 | Squa-23Ox |
| 10 | 23Ox-Lan |
| 11 | Lan-44Di |
| 12 | 44Di-14De |
| 13 | 14De-4MZC |
| 14 | 4MZC-3k4m |
| 15 | 3k4m-4MZ |
| 16 | 4MZ-Zym |
| 17 | Cholesterol production<br>(7DeC-Chol + Des-Chol) |

